# The maternal X chromosome impairs cognition and accelerates brain aging through epigenetic modulation in female mice

**DOI:** 10.1101/2022.03.09.483691

**Authors:** Samira Abdulai-Saiku, Shweta Gupta, Dan Wang, Arturo J. Moreno, Yu Huang, Deepak Srivastava, Barbara Panning, Dena B. Dubal

## Abstract

Female mammalian cells harbor two X chromosomes, one of maternal and one of paternal origin. During development, one X randomly inactivates^1–4^. This renders either the maternal or paternal X active, causing X mosaicism that varies among individual females, with some showing considerable or complete skew in the general female population^5–7^. Parent-of-X-origin can modify epigenetics via DNA methylation^8, 9^ and possibly gene expression; thus, mosaicism could buffer dysregulated processes in aging and disease. However, whether X skewing – or its mosaicism – alters functions in females is largely unknown. Here we tested whether skew toward the maternal X (Xm) influences key functions of the body. Among cardiac, bone, metabolic, and brain functions, Xm selectively impaired cognition in female mice throughout the lifespan. Cognitive deficits were accompanied by Xm-mediated acceleration of biologic or epigenetic aging of the female hippocampus, a key center for learning and memory. Xm showed epigenetic imprinting of several genes within hippocampal neurons, suggesting silenced cognitive loci. Thus, the maternal X chromosome impaired cognition, accelerated brain aging, and silenced genes. Understanding how the maternal X impairs brain function could lead to new understanding of female heterogeneity in cognitive heath and to X chromosome-derived pathways against cognitive deficits and brain aging.

## Main

Female mammalian cells harbor two X chromosomes but only one is active following embryonic development due to random X inactivation^1–4^. Thereafter, each XX cell expresses either the maternal or the paternal X chromosome, causing a cellular mosaicism in parent-of-X-origin within the organism that varies widely among females – ranging from balanced mosaicism to complete X skew^5–7^. Parent-of-X mosaicism confers both genetic and epigenetic diversity in females, which could potentially buffer dysfunction arising from processes of aging and disease. Conversely, skew toward one parental X might increase vulnerability to effects of dysregulated processes. Thus, parent-of-X origin and its skew could influence heterogeneity of health outcomes in XX individuals. Whether X skew compared to mosaicism of the active X chromosome could contribute to organismal functions in females, in the absence of X mutations, is currently unknown. We tested whether skew toward the maternal X (Xm), could impair key organ functions during middle age, when age-related dysregulation begins to arise.

We first generated mice to test whether maternal X skew (Xm) – compared to X mosaicism (Xm+Xp) – influences organ functions in aging female mice (Fig 1a). We did this by crossing mice with a targeted *Xist*-loxp deletion^10^ on the maternal X, driven by a granulosa-specific Cre^11^, which enabled germline transmission. This enforced the maternal X as the only active X chromosome. We verified mosaicism in Xm+Xp mice and paternal X silencing in Xm-only mice using immunofluorescence (Extended Data Fig. 1a, b). Since mice were on a congenic C57BL/6J background, and *Xist* is not transcribed from the active X, any differences between Xm+Xp mice and Xm mice would be attributable to epigenetic influences.

**Figure 1.**
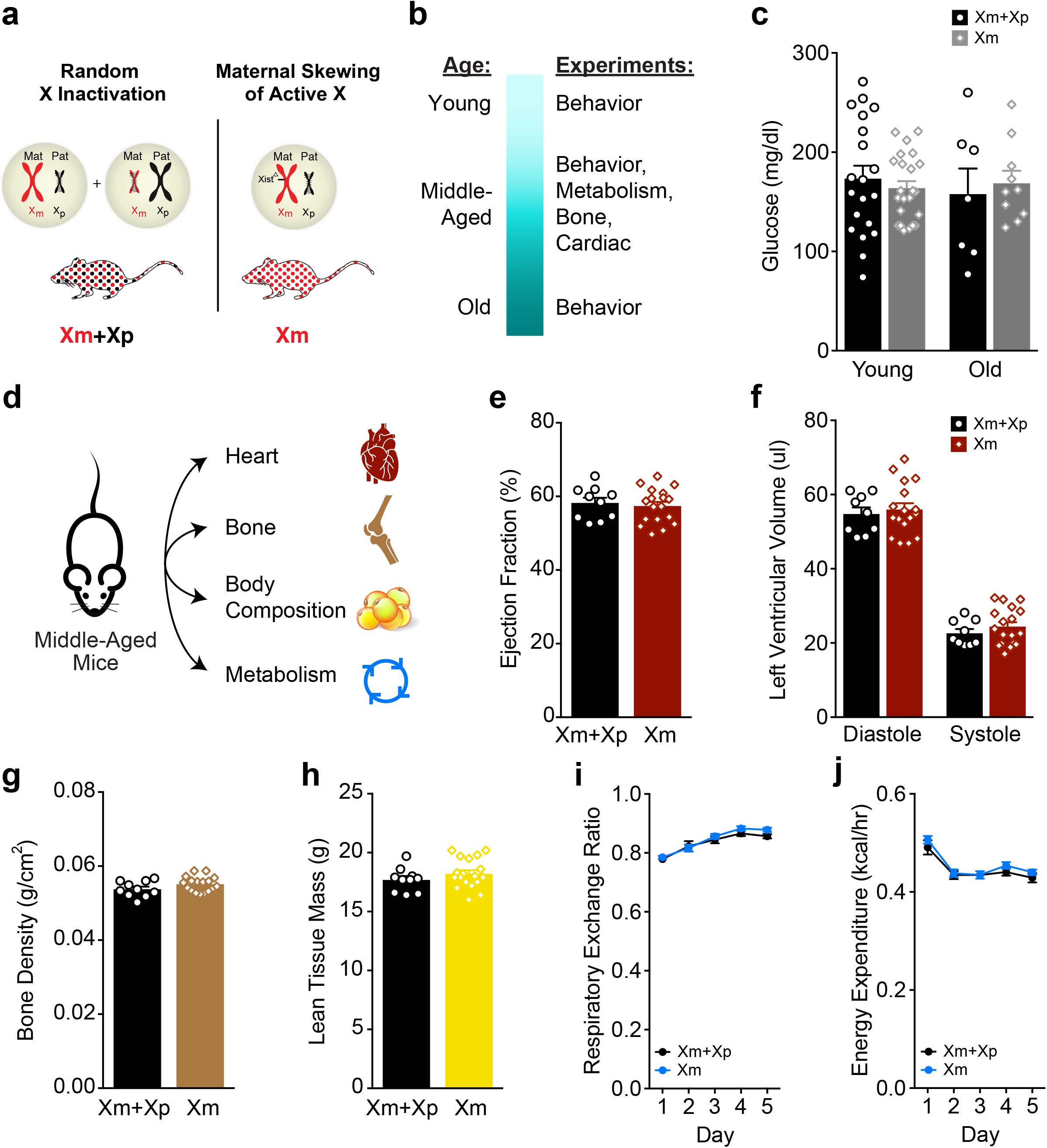
The maternal X chromosome (Xm) does not alter metabolism, body composition and cardiac function in middle-aged female mice. **a,** Diagram showing random X chromosome inactivation in wildtype, non-transgenic mice (Xm+Xp) leading to cells showing either an active paternal X (Xp) or active maternal X (Xm). In transgenic mice with maternal X skewing (Xm), all cells show only an active Xm. **b,** Experimental timeline of experiments conducted over the lifespan. **c,** Fasting blood glucose levels did not differ between Xm+Xp and Xm mice either at young (n=20-23 per experimental group; age=4-8 months) or old (n=7-10 per experimental group, age=24-27 months) ages. **d,** Diagram of organs assessed in mice during the middle-age life stage. **e, f,** Cardiac echo (n=9-18 per experimental group, age=16-19 months) was used to measure ejection fraction, and systole and diastole. **e,** Ejection fraction did not differ between Xm+Xp and Xm middle-aged mice. **f,** Left ventricular volume measurements in diastole and systole did not differ between NTG and Xm-only, middle-aged mice. **g, h,** DEXA-scan of the body (n=10-18 per experimental group, age=14-17 months) to measure bone density, and tissue mass. **g,** Bone density did not differ between Xm+Xp and Xm, middle-aged mice. **h,** Lean tissue mass did not differ between Xm+Xp and Xm, middle-aged mice. **i, j,** CLAMS metabolic cages (n=9-18 per experimental group, age=14-17 months) were used to measure metabolic parameters including respiratory exchange ratio (RER) and energy expenditure. **i,** Respiratory exchange ratio (RER) did not differ between Xm+Xp and Xm, middle-aged mice. **j,** Energy expenditure did not differ between Xm+Xp and Xm, middle-aged mice. Data represent means ± SEM.

### Maternal X chromosome and organ functions

To assess overall health, we aged littermate Xm+Xp mice and Xm mice and characterized measures across organ systems (Fig. 1b). No differences in fasting blood glucose levels were detected in young (4-8 months) or old (20-24 months) mice (Fig. 1c). We measured cardiac function, bone density, body composition, and energy metabolism (Fig. 1d) during middle-age, a life-stage vulnerable to aging-induced dysfunctions^12–16^. Echocardiography showed that ejection fraction, fractional shortening and left ventricular volume of the heart (Fig. 1e, f, Extended Data Fig. 2a) were similar between the groups. Likewise, body composition, including bone densities, lean mass and percent fat measures (Fig. 1g, h, Extended Data Fig. 2b) did not differ. Finally, oxygen consumption, carbon dioxide production, the respiratory exchange ratio, and energy expenditure were similar between groups (Fig. 1i, j, Extended Data Fig. 2c, d). Thus, maternal skewing of the active X chromosome did not alter measured heart, bone, and metabolic functions in middle-aged, female mice.

### Maternal X Chromosome Impairs Cognition

The X chromosome is enriched for genes involved in neural function^17^ and disruption of X-linked genes often causes intellectual impairments^17^. However, whether maternal X skewing, in the absence of X mutations, could influence cognition in females is unknown. Thus, we next assessed behavioral and cognitive measures in Xm (skewed) compared to Xm+Xp (mosaic) mice across the lifespan, starting with young mice (Fig. 2a).

**Figure 2.**
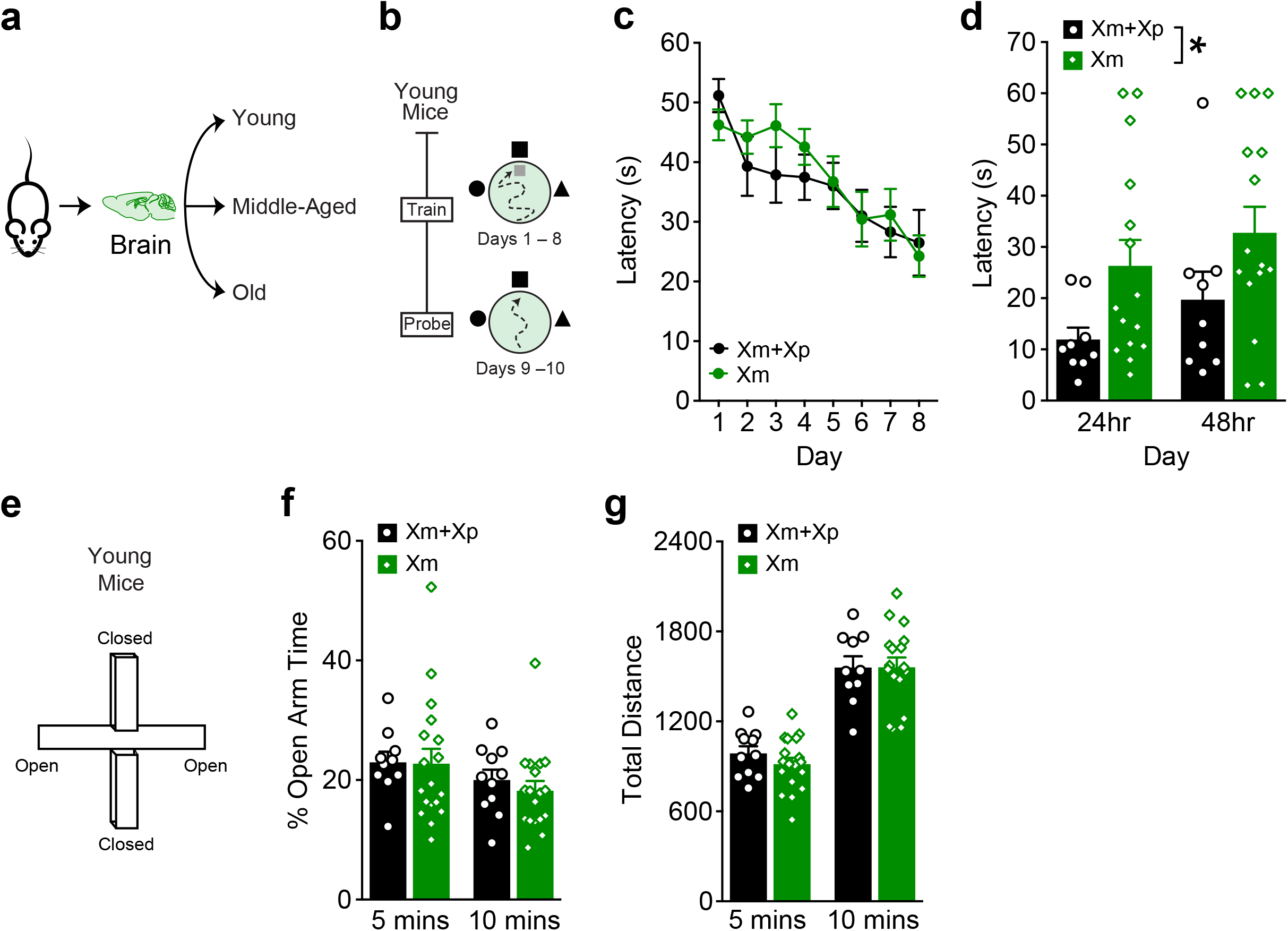
The maternal X chromosome (Xm) impairs spatial memory and learning plasticity in young female mice. **a,** Diagram of mice being tested for cognition at young, middle-age, and old life stages **b,** Paradigm for Morris water maze performed in mice (n=9-15 per experimental group, age=4-8 months). **c,** Spatial learning in the hidden trials, measured by the latency to find the platform, did not differ between the groups. **d,** Probe trials show that Xm skew impairs memory 24h and 48h after hidden training. Two-way ANOVA: genotype, \**P*<0.05. **e,** Diagram of elevated plus maze (EPM) for testing anxiety-like behavior in young (n=10-18 per experimental group, age=4-8 months) mice. **f,** Anxiety-like behavior, measured by % time in open arm, did not differ between groups at either 5 or 10 min. **g,** Total distance traveled in the EPM, did not differ between groups at either 5 or 10 min. Data represent means ± SEM.

In the Morris water maze (Fig. 2b), which measures spatial learning and memory, young (4-8 months) mice showed comparable spatial learning in trials with a hidden platform (Fig. 2c) between groups. In contrast, in probe trials, which measure ability to remember the platform location, Xm impaired memory. (Fig. 2d). Swimming speeds and ability to find a visible platform in the Morris water maze did not differ between the groups (Extended Data Fig. 3). Furthermore, time spent in open arms of the elevated plus maze (EPM), which measures anxiety-like behavior, (Fig. 2e, f), along with distance traveled in the EPM (Fig. 2g), was also similar between groups, indicating specificity of the impairments to spatial memory.

We next assessed whether maternal X skew influences another test of spatial memory, tested over the lifespan. Young, middle-aged and old Xm+Xp and Xm mice were assessed for spatial memory using repeated testing in the open field^18^ (Fig. 3a). During the young life stage, Xm increased activity at baseline followed by habituation, in the same spatial context, to similar activity levels of mosaic, Xm+Xp controls (Fig. 3b). During middle-age, Xm increased forgetfulness of the spatial context compared to mosaic, Xm+Xp controls (Fig. 3b, c). During old age, Xm further worsened forgetting (Fig. 3b,c).

**Figure 3.**
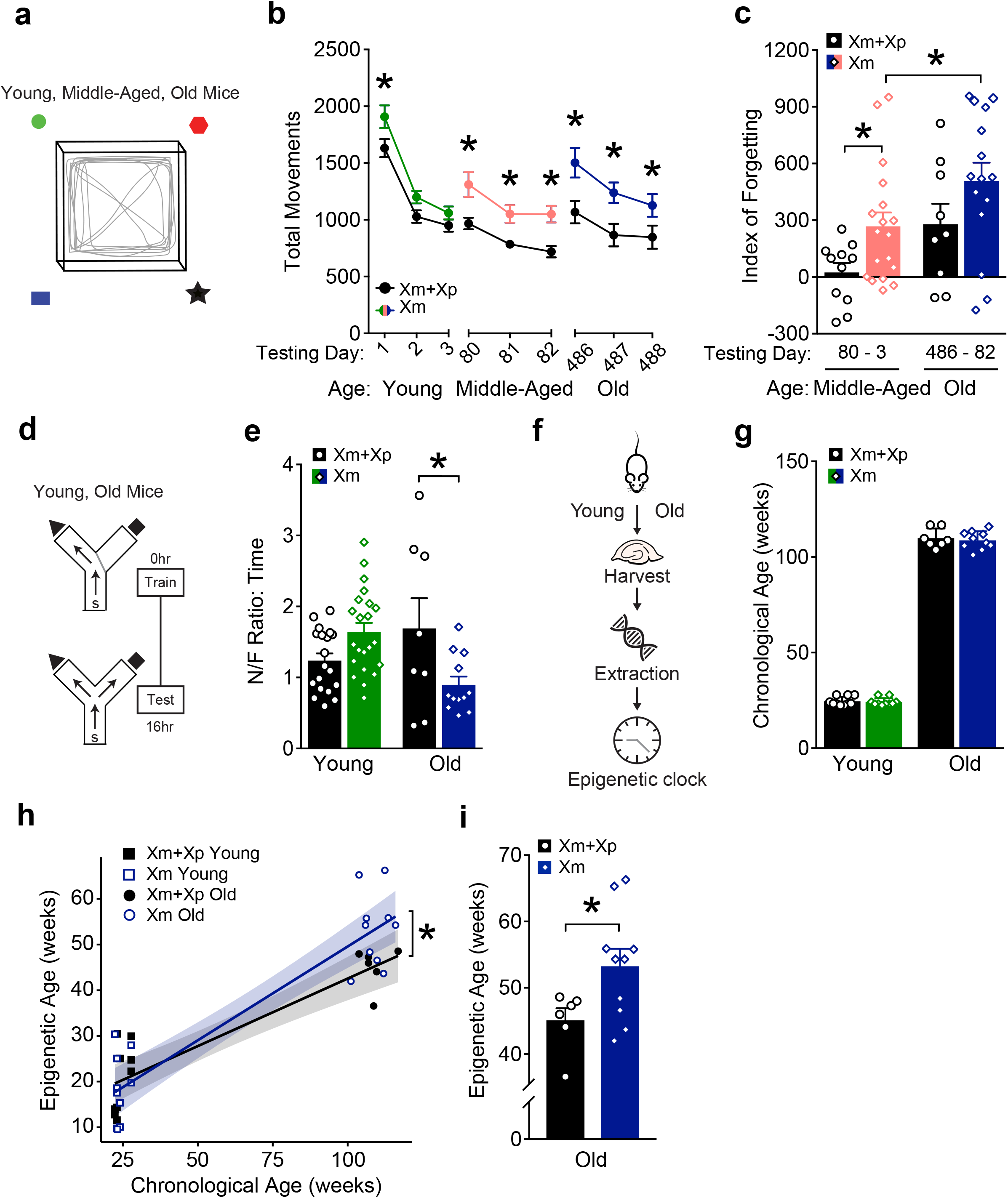
The maternal X chromosome (Xm) impairs cognition across the lifespan and accelerates epigenetic brain aging in female mice. **a,** Diagram of the open field apparatus to test context-dependent habituation and dishabituation, a test of spatial learning and memory^18^. Mice were tested for 10 minutes on 3 consecutive days across the lifespan during young (n=10-19 per experimental group, age=4-8 months), middle-age (n=10-18 per experimental group, age=9-11 months), and old (n=9-15 per experimental group, age=20-24 months) ages in the same cohort; day 1 is the first day of open field testing in young mice. **b,** Xm skew increasingly impaired spatial habituation and dishabituation across the lifespan. Two-way mixed model ANOVA: time, *P*<0.001; genotype, *P*<0.01. **c,** Xm skew accelerated age-dependent forgetfulness (measure by D80-D3) by middle-age (t-test, \**P*<0.05, Bonferroni corrected). In the old mice, Xm skew further worsened forgetting (t-test, \**P*<0.05, Bonferroni corrected). **d,** Diagram of the two-trial large Y maze for testing spatial and working memory in young (n=19-22 per experimental group, age=2-5 months) and old (n=8-12 per experimental group, age=20-24 months) mice, tested simultaneously. **e,** In old, but not young mice, Xm skew impaired spatial and working memory, measured by ratio of time spent in the novel/familiar arm. Two-way ANOVA: age by genotype interaction; *P*<0.01. \**P*<0.05 (Bonferroni-corrected) **f,** Diagram of experimental process for assaying epigenetic, DNA age in young (age=6 months, n=10 per experimental group) and old (age=25 months, n=6-10 per experimental group) mice. **g,** Chronological age of young mice or old mice did not differ by experimental group. **h,** Epigenetic age plotted against chronological age shows Xm skew accelerated biologic aging after the young life stage (\**P*=0.05, Linear regression analysis). **i,** Xm skew increased epigenetic age of the hippocampus in the old life-stage \**P*<0.05 (t-test).

Xm increasingly worsened memory with age; thus, we next tested young and old mice of each group in the two-trial Y maze (Fig. 3d), a task sensitive to deficits in working and spatial memory in aging^19, 20^. During the young lifestage, measures did not differ between Xm and Xm+Xp mice (Fig. 3e). During old age, Xm decreased the ratio of time mice spent in the novel compared to familiar arm, indicating worsened memory compared to mosaic Xm+Xp controls (Fig. 3e). Thus, Xm impaired working and spatial memory in old female mice.

### Accelerated biological aging by the maternal X chromosome

Since the maternal X chromosome consistently worsened cognitive dysfunction in aging, we wondered whether it accelerates biological aging of the hippocampus. Aging induces epigenetic alterations that are robust indicators of biological age^21^, measured as predictable DNA methylation (DNAme) patterns^22, 23^, and termed the “epigenetic clock”. Acceleration of the epigenetic clock in one experimental group compared to another indicates increased biological aging. To assess the relative DNAme of Xm compared to Xm+Xp hippocampi, we analyzed methylation profiles from >500 specific age-associated loci of DNA in young and old mice in each experimental group (Fig. 3f). Chronological ages between the experimental groups did not differ (Fig. 3g). In contrast, biological ages differed; Xm accelerated the epigenetic clock compared to mosaic, Xm+Xp controls (Fig. 3h), causing Xm hippocampi to be biologically older (Fig. 3i).

Among key cardiac, bone, metabolic, and brain functions, the maternal X chromosome selectively impaired brain function. Several lines of evidence support disproportionate X chromosome influence on the brain. Disruptions in X gene expression, through X-linked disorders, often cause intellectual disability^17, 24^. Furthermore, in the brain, more genes are expressed from the X chromosome than from any other single autosome^25^. Together, these examples imply that the brain, compared to other organs, may be more sensitive to variations in X chromosome expression.

In our study, the maternal and paternal X chromosomes were genetically identical; thus, maternal X chromosome skew would cause epigenetic differences that influence gene expression. Among these epigenetic differences, effects of *Xist* deletion on the active X, which enforced Xm skew, cannot be ruled out, though it is normally silenced on the active X. Studies of female individuals with Turner’s syndrome that harbor only a maternal, compared to a paternal, X chromosome show cognitive deficits^26^. These human data suggest that genes influencing cognition are imprinted, or silenced, on the maternal X; however, whether Xm undergoes imprinting, or gene silencing, remains unknown.

### Epigenetic silencing by the maternal X chromosome

We next investigated whether the maternal X chromosome undergoes gene silencing, an epigenetic parent-of-origin effect. To achieve high resolution in this study, we applied deep RNA sequencing to transgenic female mice with nuclear localized genetic reporters that enabled neuron-specific identification of Xm (labeled by GFP) and Xp (labeled by tdTomato) cells^27^. Using cell sorting, we separated Xm from Xp neurons from female XX hippocampi that underwent random X chromosome inactivation and conducted informatics (Fig. 4a,b). We detected 848 of approximately 1500 known X chromosome genes and applied established criteria to detect imprinting^28, 29^. The maternal X chromosome showed silencing, or imprinting, of 9 genes as shown in the heatmap (Fig. 4c), including *Sash 3, Tlr13, Tlr7,* and *Cysltr,* the most robustly silenced genes. Genes were distributed throughout the X chromosome (Fig. 4d) and nearly undetectable from the maternal X, with very high expression from the paternal X (Fig. 4e-h). Additionally, the paternal X chromosome showed silencing of 2 genes, *Xlr3b* and *Trpc5* (Fig. 4c). *Xlr3b,* as previously identified^30, 31^, was nearly undetectable from the paternal X chromosome, and showed very high expression on the maternal X chromosome (Fig. 4i). We further validated our RNA-seq data by qRT-PCR of Sash 3, Tlr13, Tlr7, Cysltr1and Xlr3b mRNA (Fig. 4j) and found similar expression patterns (Fig. 4k-o).

**Figure 4.**
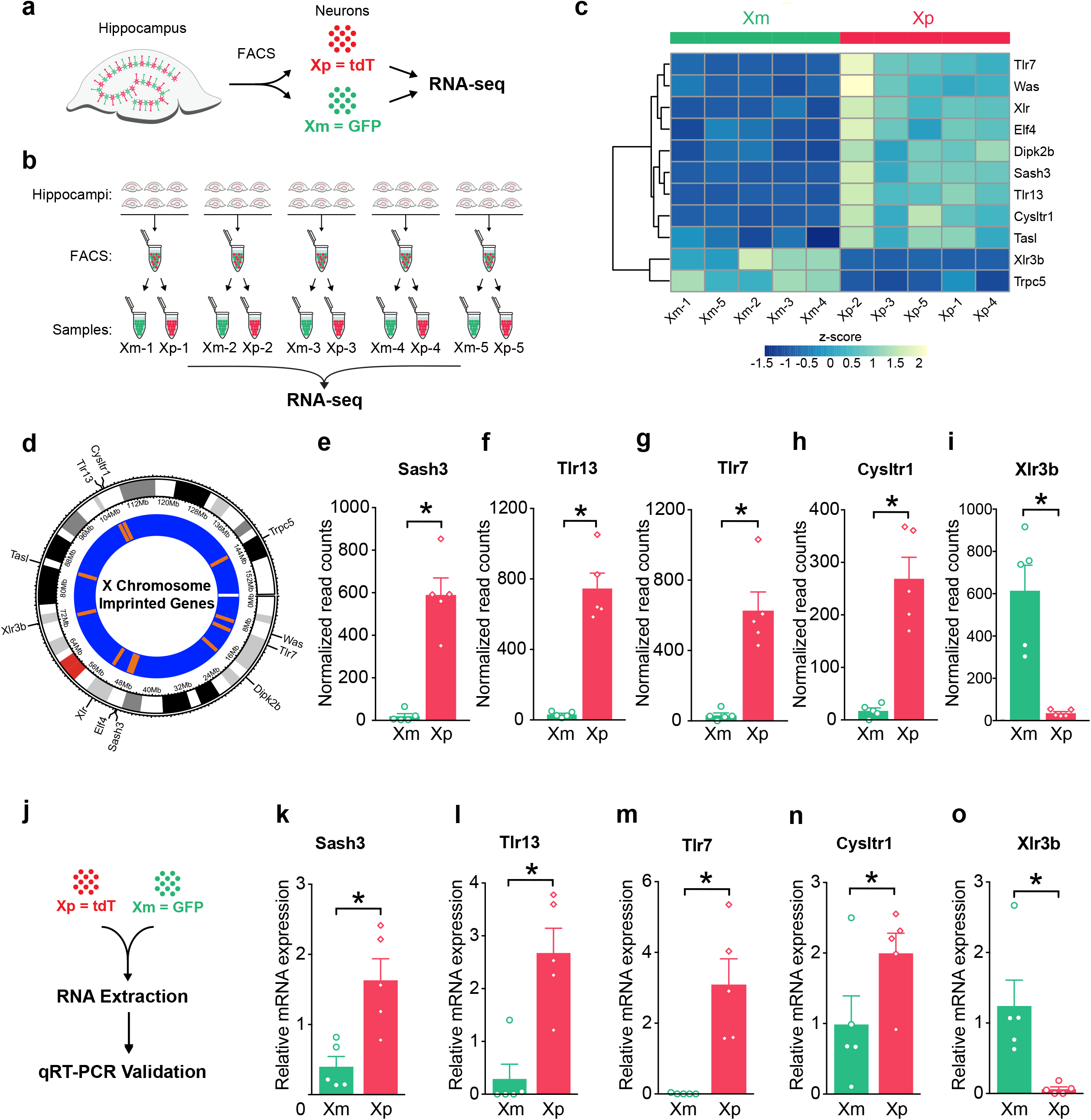
The identification of imprinted X genes on the maternal X (Xm) and paternal X (Xp) chromosome of female hippocampal neurons. **a,** Diagram of experimental paradigm. Neurons labeled with synapsin 1 driven by Cre were FACS-sorted into Xp (paternal X origin, tdT red) or Xm (maternal X origin, GFP) from the same hippocampus, for RNA sequencing. **b**, Hippocampi of adult female mice underwent sorting to obtain Xm- and Xp-expressing neurons (age=3-4 months; n=5 samples per each group; each sample contained cells from 6 pooled hippocampi) for RNA isolation for RNAseq and for qRT-PCR validation. **c,** Heatmap of gene expression for top 11 imprinted genes on the X chromosome. **d,** Circos plot showing topographical distribution of imprinted genes on the X chromosome. **e-I,** RNA-seq expression graphs for top 5 most robustly X-imprinted genes including (**e**) Sash3, (**f**) Tlr13, (g), Tlr7, (**h**) Cysltr1, (**z**), Xlr3b (adjusted *P* values, DESeq2 analysis: \**P*<0.05). **j,** Diagram of qRT-PCR experimental workflow for validation of RNAseq results. RNA samples were derived from samples outlined in (b) (female mice, age=3-4 months; n=5 samples per each group; each sample contained cells from 6 pooled hippocampi). **k-o,** qPCR validation for the top 5 imprinted genes including (**k**) Sash3, (**l**) Tlr13, (**m**) Tlr7, (**n**) Cysltr1 (**o**) Xlr3b (Paired one-tailed t-tests: \**P*<0.05).

## Discussion

We found that skew toward the maternal X chromosome, compared to X mosaicism, impaired cognition across the lifespan and accelerated brain aging. Xm and Xp were identical in our studies, suggesting an epigenetic, parent-of-origin modulation of X chromosome gene expression. Indeed, Xm neurons underwent silencing of select genes, when directly compared to neighboring Xp neurons, representing maternal imprinting of the female hippocampus, a key center for memory.

Maternal X skewing selectively impaired brain functions, among multiple organ systems studied in normal aging females – and this impairment was observed across the lifespan. In the brain, more genes are expressed from the X chromosome than from any single autosome^32, 33^; thus, the brain is probably more sensitive to alterations of X gene expression. In the context of X chromosomal abnormalities, female individuals with Turner’s syndrome (45, XO) experience greater cognitive impairment when the X is maternally, compared to paternally, inherited^17, 34^, a finding also observed in Turner’s mouse models^30, 35^. However, even in the absence of mutations or X chromosomal abnormalities, our data suggest that the maternal X, which is variably skewed in the population of typical females with two X’s, impairs neural functions.

We investigated X imprinting in a select population of hippocampal cells that express synapsin 1, a neuronal and synaptic marker^36^. Assessing this subtype of hippocampal cells increases the relevance of X imprinting to mechanisms of learning and memory and represents an advance from historic assessments of whole brain or regional homogenates that necessarily amassed together multiple cell types. Our data in neuronal hippocampus cells indicate maternal X silencing of possible cognition-related loci.

The majority of X imprinting in hippocampal neurons was of maternal origin. Notably, our data validate *Xlr3b* as a known paternally imprinted gene, as identified in previous screens^30, 31^. Among the top maternally imprinted genes, *Tlr7* regulates genes crucial for memory and long-term potentiation (LTP)^37, 38^. All of the top maternally silenced genes *Tlr7, Sash 3, Cysltr,* and *Tlr13* are involved in immune-related processes - and their roles in neuronal functions have largely been unexplored. It is interesting to speculate a role for these factors at the intersection between immune signaling and synaptic pruning of neurons by microglia^39^, critical to optimal synaptic connectivity. The silencing of these X factors in Xm neurons, compared to their robust expression in Xp neurons, may impair substrates and signals of cognition.

Our data suggest that females with more skew toward the maternal X chromosome, even in the absence of mutations, could experience decreased cognitive functions – or be at increased risk for neurodegenerative disease – compared to those with more balanced X mosaicism in parent-of-X origin, particularly in aging. This may be due to the absence of certain X genes in hippocampal neurons expressing the maternal X. Understanding epigenetic parent-of-X silencing in neurons could lead to unraveling X chromosome-derived pathways that can counter cognitive deficits and brain aging.

## MATERIALS AND METHODS

### Animals

The Institutional Animal Care and Use Committee of the University of California San Francisco approved all animal studies. Mice were kept on a 12 hr light/ dark cycle with *ad libitum* access to food and water. The standard housing conditions were 5 mice per cage except during the Morris Water Maze and the CLAMS metabolic tests when mice were single-housed. All experiments were carried out during the light cycle with the exception of the CLAMS metabolism test that collected data throughout the light and dark cycles. To assess effects of Xm skew compared on mosaicism, we generated mice with global maternal skewing of the active X chromosome (Xm) with non-transgenic littermates showing normal, random X inactivation (Xm+Xp). Maternal skewing of the active X was achieved via *Xist* deletion. Of note, Xist is a long noncoding RNA that regulates random X chromosome inactivation. Briefly, we crossed 129-*Xist^tm2Jae^*/Mmnc mice^10^ obtained from the Mutant Mouse Resource and Research Centers (MMRRC) with Zp3-Cre mice, generously provided by Sundeep Kalantry^11^. F2 mice were then backcrossed to C57BL/6J to obtain a congenic C57BL/6J background, verified by genetic testing. Female mice underwent multiple tests of metabolism, cardiac function, body composition, behavior and cognition during life stages indicated in figure legends. All arenas and equipment were cleaned with 70% ethanol between tests, except for the water maze. All experimenters were blind to mice genotypes and groups.

To assess the differences between Xm and Xp neurons from the same brain, we used well-characterized mice^27^ generated to carry X-linked, Cre-activated, and nuclear-localized fluorescent reporters of GFP on one X chromosome and tdTomato on the other. They were also backcrossed to obtain a congenic C57BL/6J background. These reporter mice possess a floxed tdTomato fluorescence protein or a floxed GFP fluorescence protein inserted into the *Hprt* locus of the X chromosome using modified *Hprt* targeting vectors. The *Hprt* locus is subject to random XCI and inserting the fluorescent proteins in this position ensures that once crossed with a suitable Cre line, either GFP or tdTomato is expressed from each cell but never both. The Cre line used to drive cell-type specific Xm and Xp fluorescence was with a synapsin I, neuron-specific promoter, well characterized^36^.

### Cardiac Function

Cardiac function measurements were performed as described^40^. Briefly, mice were anaesthetized with isoflurane. Body temperature was monitored throughout the procedure using a rectal probe. A warm ultrasound gel was applied to the chest. Using MX550S transducer, the B and M mode parasternal short-axis view was recorded, the diameter of the left ventricular lumen was measured, then ejection fraction was calculated. Afterwards, electrodes were removed, ultrasound gel was cleaned, and animals were allowed to recover before being returned to their cages.

### Body Composition Analysis

Body composition analysis was conducted as described^41, 42^ in conjunction with the metabolism core at the Nutrition and Obesity Research Center at UCSF. The Lunar PIXImus densitometer (GE Medical Systems, Madison, WI) was used to obtain body composition analysis of each mouse using dual energy x-ray absorptiometry (DEXA) technology. Briefly, mice were weighed and anaesthetized with avertin before being immobilized on a sticky mat. X-ray measurements were taken, and the region of interest was adjusted to ensure the whole mouse was considered in the analysis. Data output provided bone, tissue and fat measurements.

### CLAMS Metabolism

Metabolic analysis was conducted as previously described^41^ with the metabolism core at the Nutrition and Obesity Research Center at UCSF. Briefly, mice were single-housed for one week for habituation to the experimental conditions. Mice were then placed in the Comprehensive Lab Animal Monitoring System (CLAMS) and monitored over five days. Data was generated in 1 hr bins and used to calculate metabolic parameters including oxygen consumption (VO2), carbon dioxide production (VCO2), energy expenditure (EE) and respiratory exchange ratio (RER).

### Morris Water Maze

Water maze testing was performed as described^43–46^. Briefly, we filled the water maze pool (diameter, 122 cm) with white opaque water (21° ± 1°C) and submerged a square 14 cm^2^ platform 2 cm below the surface. Mice underwent two pretraining trials that consisted of swimming through a channel to mount a rescue platform, prior to hidden training. The platform was kept in the same, submerged spot during all hidden platform training trials; the drop location varied between trials. For the hidden trials, mice received 4 trials daily for 8 days. For the probe trials, the platform was removed, mice were allowed to swim for 60 seconds, and their latency to enter the previous platform area was recorded. Following probe testing, mice were assessed for their ability to find the platform when it is marked with a visible cue (15 cm pole on the platform).

### Open Field

Open field testing was carried out as described^18^. Briefly, mice were acclimated to the room for 1 hour before testing and allowed to explore the open field for 10 minutes. The open field consisted of a clear plastic chamber (41 x 30 cm) and total activity was detected by measuring beam breaks using an automated Flex-Field/Open Field Photobeam Activity System (San Diego Instruments).

### Elevated Plus Maze

Elevated Plus Maze testing was carried out as described^43^. The room was maintained in dim light for both habituation and testing. Briefly, mice were habituated to the testing room for 1 hour before testing. Mice were placed in the center of the elevated plus maze facing the open arm and allowed to explore for 10 min. Distance traveled and % time spent in the open versus closed arms was recorded using the Kinder Scientific Elevated Plus Maze and MotorMonitor™ system.

### Large Y Maze

The large Y maze two-trial test (with visual cues at the end of each arm) was carried out as described^45, 47^. Sixteen hours after a training session during which the novel arm was closed off, mice were returned to the large Y maze and allowed to explore all arms freely for 5 minutes. Time spent in the novel and familiar arm was recorded using the AnyMaze software and the novel to familiar ratio was calculated.

### Epigenetic DNA age analysis (DNAme age)

Hippocampal tissue samples were flash-frozen. Samples then underwent sample library preparation and sequencing analysis as described (Zymo Research)^48^. Briefly, genomic DNA was extracted using the Quick-DNA Miniprep plus kit and bisulfite converted using the EZ DNA Methylation Lightning kit. The samples were then enriched for sequencing of >500 age-associated gene loci on Illumina HiSeq instrument. Illumina’s base calling software was used to identify sequence reads and aligned to a reference genome using Bismark, an aligner optimized for bisulphite sequence calling (http://www.bioinformatics.babraham.ac.uk/projects/bismark/). The methylation level was determined by proportion of the numbers of ‘C’ reported to the total numbers of ‘C’ and ‘T’. Calculated DNA methylation values obtained from the sequence data were used for epigenetic age prediction using a proprietary DNAge predictor (Zymo).

### RNA-seq

RNA-seq was conducted on hippocampal cells separated into maternal X active (Xm) and paternal X active (Xp) cells. Hippocampi from 6 mice were pulled together to form a single sample. In total, 5 samples derived from 30 mice were prepared. Each sample was FACS-sorted into Xm (GFP+, green) cells and Xp (tdT+, red) cells. The samples were stored in TRIzol and then underwent RNA-seq analysis. Briefly, RNA sequencing libraries were prepared using the SMART-Seq v4Ultra Low Input RNA Kit (Clonetch). Paired-end reads are obtained from an Illumina Hi-Seq instrument. Quality of reads was determined using FastQC, and more than 90 percent of reads from each sample had a mean quality score >30. The trimmed reads were mapped to the *Mus musculus* GRCm38 reference genome available on ENSEMBL using the STAR aligner v.2.5.2b. Unique gene hit counts from exons were calculated using featureCounts from the Subread package v.1.5.2. Downstream differential expression analysis was performed using DESeq2.

### Identification of Imprinted Genes and qPCR Validation

Imprinted genes were selected based on a list of criteria as follows: 1) Significant *p* value, and adjusted *p* value. 2) Sum of normalized gene expression from one sample group <50 and sum of normalized gene expression from the other sample group >100. 3) Significant Chi-square *p* value, and adjusted significant Chi-square *p* value, and 4) Fold change >10. Based on these criteria, we identified 5 imprinted genes. RT-qPCR was used to validate the expression of imprinted genes identified from the RNA-seq analysis. Primers were designed using the NCBI Primer Blast page and purchased from IDT. *Sash3* Fwd: CTGGCAGTGAAGAGGCTGAA, Rev: GACCCTGCAGTTGCTCTTCT; *Cysltr1* Fwd: GGTACCAGATAGAGGTCTCCC, Rev: CTCCAGGAATGTCTGCTTGGT; *Tlr13* Fwd:TCCTCCCTCCCTGGAGTTTT, Rev: AGGCACCTTCGTCGATCTTC; *Tlr7* Fwd: TGCACTCTTCGCAGCAACTA, Rev: ATGTCTCTTGCTGCCCCAAA; *Xlr3b* Fwd: AAAAGGAAGGCCACTGACAC, Rev: ACCAGCATCAAGGACTTCTCTG; *GAPDH* Fwd: GGGAAGCCCATCACCATCTT, Rev: GCCTTCTCCATGGTGGTGAA; *18sRNA* Fwd: AGGGGAGAGCGGGTAAGAGA, Rev: GGACAGGACTAGGCGGAACA.

### Statistical Analysis

Experimenters were blinded to genotype. Statistical analyses were carried out using GraphPad Prism (version 7.0) for t-tests and two-way ANOVAs and R Studio (v 2.0) for post hoc tests. All tests were two-tailed unless indicated otherwise. Differences between two means was assessed using unpaired t-tests and a two-way ANOVA to assess differences among multiple means for all experiments unless otherwise stated. Post hoc tests were conducted with Bonferroni-Holm correction to control for a family-wise error rate at α=0.05. A mixed model ANOVA was used to analyze Morris water maze and open field data and included effects of repeated measures. Error bars represent SEM and null hypotheses were rejected at or below a *P* value of 0.05. Linear regression analysis to compare the chronological and epigenetic ages was done in R. Linear models were fit in R using the standard lme package.

## Acknowledgements

We thank Sundeep Kalantry for providing mice with an *Xist* deletion and Chen Chen for mouse colony management. FACS analysis was supported in part by Helen Diller Family Comprehensive Cancer Center (HDFCCC) Laboratory for Cell Analysis Shared Resource Facility through NIH (P30CA082103). Metabolic analyses were supported in part by the Nutrition and Obesity Research Center (NORC) Mouse Metabolism Core through NIH (P30DK098722). Primary funding for the study was by NIH grants NS092918 (D.B.D.) and AG068325 (D.B.D.), American Federation for Aging Research (AFAR)(D.B.D.), and philanthropy (D.B.D.).

## Author contributions

S.A.S. and D.B.D. conceived and designed all experiments. S.A.S. and D.W. performed all behavioral experiments. Y.H. and D.S. performed cardiac measurement experiments. S.A.S. performed RNA-seq and validation experiments with input from B.P. S.A.S., and S.G. analyzed transcriptomic data. S.A.S, A.M. and D.B.D. analyzed all the remaining data in the manuscript. S.A.S and D.B.D. wrote this manuscript with input from B.P.

