## Supplementary Figures for "The maternal X chromosome impairs cognition and accelerates brain aging through epigenetic modulation in female mice"

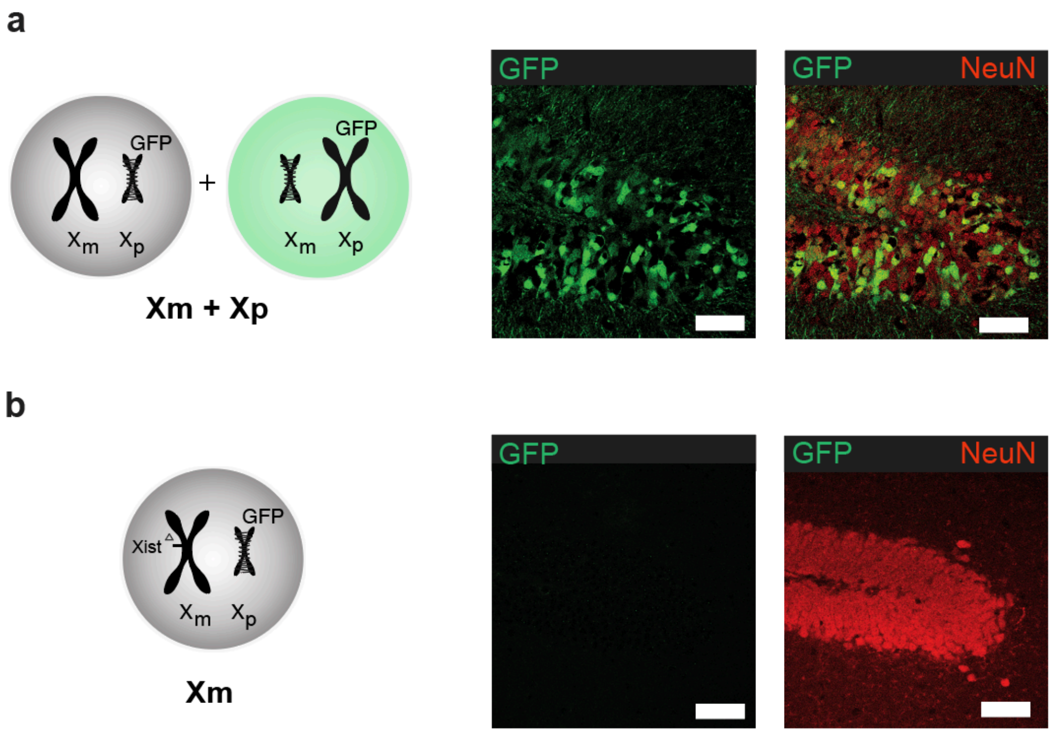


**Supplementary Figure 1. Parent-of-X mosaicism compared with paternal X silencing in maternal X skew in the hippocampus of a female mouse.**

**a**, Left, diagram of parent-of-X composition of cells in Xm+Xp mice. Right, Representative image of immunohistochemistry showing cells that express Xp (tagged with GFP) and all NeuN+ cells (RFP) in the dentate gyrus region of the hippocampus in a female mouse. Scale Bar=60μm.

**b,** Left, diagram of cellular composition of cells in which the paternal X is silenced and skewed toward Xm (due to Xist deletion on Xm). Right, Representative image of immunohistochemistry showing maternal X skew, via paternal silencing of X (Xp, tagged with GFP) cells in the dentate gyrus region of the female hippocampus. All neurons are tagged with NeuN (red). Scale Bar=60μm.

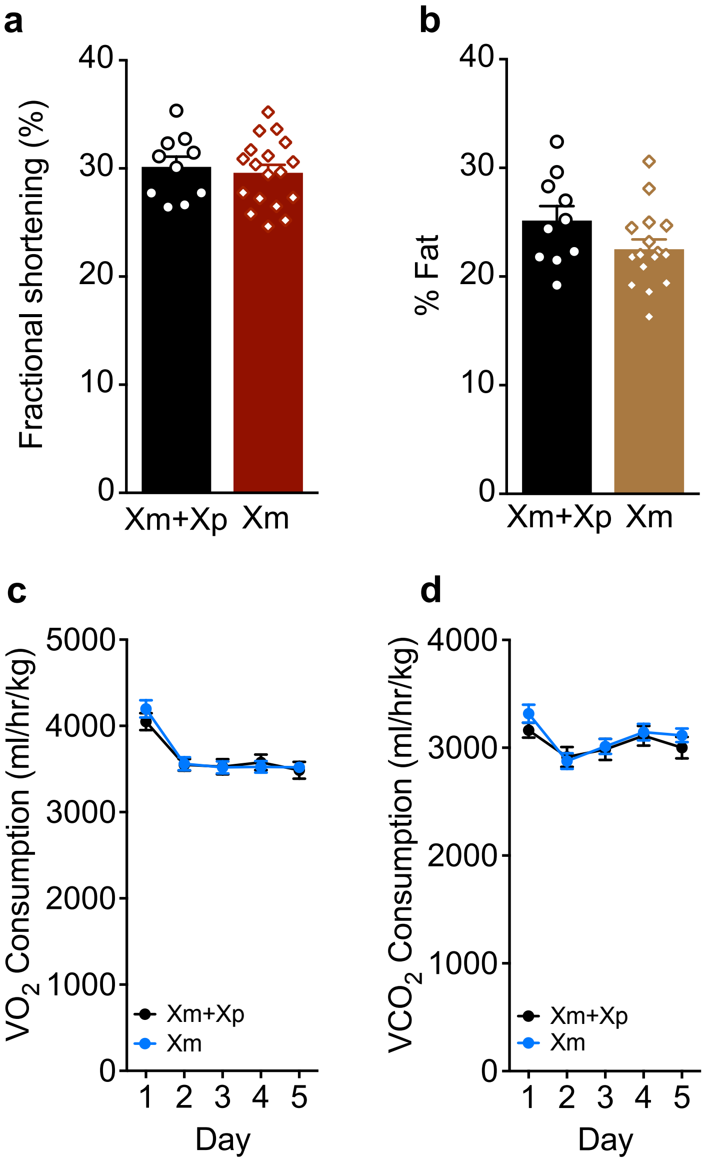


**Supplementary Figure 2. Additional measures in the body including cardiac, % fat, and V02 and VCO2 did not differ between experimental groups**.

**a.** Cardiac echo measurements showed equivalent measures between Xm+Xp and Xm mice in fractional shortening (age=14-17 months, n=10-18 per experimental group). Unpaired two-tailed t-test, *P*>0.1**.**

**b.** Body fat percentage did not differ between Xm+Xp and Xm mice (age=14-17 months, n=9-17 per experimental group). Unpaired two-tailed t-test, $P>0.05$**.**

**c, d.** The CLAMS metabolic cages were used to measure different metabolic parameters including oxygen consumption (VO2), and carbon dioxide production (VCO2) (age=14-17 months, n=10-18 per experimental group).

**c,** VO2 consumption did not differ between the groups. Two-way ANOVA: genotype $P>0.05$

**d,** CO2 consumption did not differ between the groups. Two-way ANOVA: genotype $P>0.05$

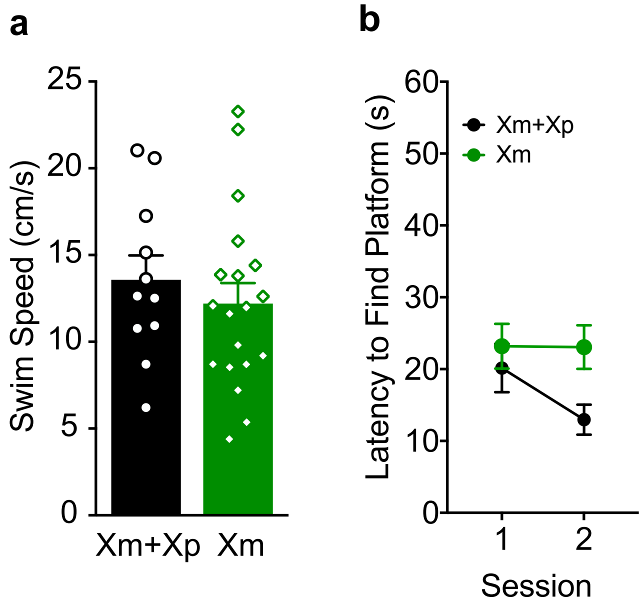


**Supplementary Figure 3. Swim speed and latency to find visible platform did not differ between groups.**

**a,** Swim speed did not differ between the experimental groups, measured during the visible platform trials (age=4-8 months, n=11-19 per experimental group). Unpaired two-tailed t-test, $P>0.05$**.**

**b,** Latency to find a visible platform did not differ between the experimental groups. (age=4-8 months, n=11-19 per experimental group). Mixed model ANOVA: genotype $P>0.05$.
